# Seeing cells smell: Dynamic optical measurements of Ca^2+^ and cAMP signaling from Olfactory Receptors transiently expressed in HEK293TN cells

**DOI:** 10.1101/771261

**Authors:** A Pietraszewska-Bogiel, L van Weeren, J Goedhart

## Abstract

Olfactory receptors (ORs) constitute the largest family of G-protein coupled receptors. They are responsible for the perception of odor (olfaction) and also play important roles in other biological processes, including regulation of cell proliferation. Their increasing diagnostic and therapeutic potential, especially for cancer research, requests the ongoing development of methodologies that would allow their robust functional expression in non-olfactory cells, and dynamic analysis of their signaling pathways. To enable realtime detection of OR activity, we use single cell imaging with genetically encoded fluorescent biosensors, Yellow Cameleon or EPAC, which are routinely used for kinetic measurements of Ca^2+^ or cAMP signaling downstream of various G-protein coupled receptors. We demonstrate that the co-expression of Lucy-Rho tagged variants of ORs together with an accessory protein, RTP1s, in HEK293TN cells is sufficient to detect the activity of a panel of ORs. Using this methodology, we were able to detect both Ca^2+^ and cAMP signaling downstream of twelve ORs within 2 minutes from the application of odorant.

## INTRODUCTION

Olfactory (or odorant) receptors (ORs) are G-protein coupled receptors (GPCRs) expressed at the cell surface of olfactory sensory neurons (OSNs) in the main olfactory epithelium (OE) (Buck & Axel 1991, Verbeurgt et al. 2014, Olender et al. 2016). They detect exogenous chemical ligands, referred to as odorants, and serve as the chemical sensors of smell. The odorant signal transduction is initiated when odorants interact with specific ORs, resulting in the OR coupling to the olfactory G protein (an Gα_olf_/Gß_1_/Gγ_13_ heterotrimer; reviewed in Su et al. 2009 and Li et al. 2013a). This results in an activation of type III adenylyl cyclase (ACIII), followed by the generation of cAMP and opening of the cation-selective cyclic nucleotide-gated channels. The resulting influx of extracellular Ca^2+^ leads to depolarization of OSNs, which fire action potentials to the brain (for the role of Cl^−^ outflow through a calcium-activated Cl^−^ channel in olfaction see Dibattista et al. 2017). ORs are also required for a proper targeting of olfactory neuron axons to their corresponding glomeruli in the olfactory bulb (Feinstein et al. 2004, Richard et al. 2013), and some ORs have recently been shown to underlie the response to both chemical and mechanical stimuli in OSNs (Connelly et al. 2015). But it is not all they do. Despite their name, many human OR genes are expressed in non-olfactory tissues (Feldmesser et al. 2006, Flegel et al. 2013, Zhang et al. 2004, 2007) and have been implicated in: cytokinesis, cell proliferation and migration (Neuhaus et al. 2009, Zhang et al.2012, Busse et al.2014, Sanz et al. 2014, Gelis et al. 2016, Manteniotis et al. 2016ab, Jovancevic et al. 2017a, Tsai et al. 2017, Weber et al. 2017), sperm and monocyte chemotaxis (Spehr et al. 2003, 2006, Veitinger et al. 2011, Li et al. 2013b), melanocyte function (Gelis et al. 2016, 2017), serotonin release (Braun et al. 2007, Kidd et al. 2008, Gu et al. 2014, Kalbe et al. 2016), insulin secretion (Munakata et al. 2018), myocardial function (Kim et al. 2015, Jovancevic et al. 2017c), and regulation of breathing (Chang et al. 2015).

Humans have more than 300 intact ORs (Malnic et al. 2004) but odorant ligands have been published only for approximately 50 receptors (reviewed in Mainland et al. 2014 and Lunde et al. 2012, Jaeger et al. 2013, Busse et al. 2014, Shirasu et al. 2014, Gonzalez-Kristeller et al. 2015, Manteniotis et al. 2016ab, Weber et al. 2017). This is due in part to difficulties with functionally expressing ORs in heterologous systems, such as human embryonic kidney (HEK) 293 cells, routinely used for throughput analyses of other GPCRs. Olfactory neurons have a selective molecular machinery that promotes proper targeting of ORs to the cell surface. In contrast, ORs expressed in heterologous cell systems are most often retained in the ER, resulting in OR degradation (Lu et al. 2003). Optimal cell surface expression often requires their co-expression with OSN specific accessory proteins (Receptor Transporting Protein-1 and -2, RTP1 and RTP2; Receptor Expression Enhancing Protein-1, REEP1) or elements of canonical olfactory signaling pathway (Gα_olf_ and guanine nucleotide exchange factor, Ric8B), and modification of the OR itself (most often by stably fusing the first 20 amino acids of rhodopsin, so called Rho tag, to the receptor’s N terminus) (Krautwurst et al. 1998, Saito et al. 2004, 2009, Von Dannecker et al. 2005, 2006, Neuhaus et al. 2006, Kerr et al. 2008, Zhuang & Matsunami 2007, Wu et al. 2012, Yu et al. 2017, reviewed in Peterlin et al. 2014). Recently, a cleavable 17-amino acid leucine-rich N-terminal signal peptide (Lucy tag) was shown to promote cell surface OR expression in HEK293 cells (Shepard et al. 2013).

Moreover, the functional readout employed for most odorant screening analyses involves Gα_olf_-dependent activation of endogenously expressed adenylyl cyclase. The resulting increase in cAMP is most often monitored via a cAMP response element-driven luciferase reporter and the resulting light production (Saito et al. 2004). However, a significant drawback of this system is the time required to achieve measurable and consistent odorant-dependent luciferase production (up to four hours of odorant stimulation on a standard luminometer plate reader). The prolonged exposure to odorant in these experiments may lead to adaptation.

In view of the increasing interest in odorant receptors (fueled by their emerging diagnostic and therapeutic potential in humans), we evaluated some of the strategies available for functional analyses of ORs. Our aim was to record OR activity in real time and in a system that would require co-expression of a minimal number of additional constructs.

## RESULTS and DISCUSSION

### OR-mediated Ca^2+^-transients and cAMP production in HEK293TN cells occur almost instantaneously after the odorant addition

We used a subset of twelve human ORs with previously identified ligands (Table 1). We expressed Rho tagged (abbreviated as RhoOR) and Lucy-Rho tagged (abbreviated as LROR) variants of these ORs in HEK293TN cells together with a biosensor and a short form of RTP1 (RTP1s), as it was shown to have more robust effect on promotion of OR trafficking and odorant-induced response of ORs than RTP1, RTP2 or REEP1 (Zhuang & Matsunami 2007). Moreover, OR-mediated signaling in heterologous systems can vary from the canonical (Gα_olf_-ACIII-Ca^2+^ influx) olfactory signaling pathway operating in OSNs (Shirokova et al. 2005, Oka et al. 2006, Hamana et al. 2010). Therefore, we have chosen to monitor possible changes in both intracellular Ca^2+^ and cAMP levels in odorant-stimulated cells with the Ca^2+^-biosensor, Yellow Cameleon (YC3.6, Nagai et al. 2004), and the cAMP biosensor, EPAC (^T^EPAC^VV^; Klarenbeek et al. 2011).

**Table 1.** Summary of OR-odorant pairs analyzed, including the odorant-induced endogenous response in HEK293TN cells. From left to right: name of the OR tested and its expression (as reported previously); odorant and concentration tested; odorant-mediated, integrated YC3.6 signals of HEK293TN cells expressing RTP1s, biosensor, and: no additional OR, RhoOR construct or LROR construct; odorant-mediated, integrated EPAC signals of HEK293TN cells expressing RTP1s, biosensor, and: no additional OR, RhoOR construct or LROR construct. Integrated values for YC3.6 and EPAC signals were obtained by summing all values in the 38–158 s time window and dividing them by the number of cells. YC3.6 and EPAC signals in control HEK293TN cells stimulated with DMSO are given at the bottom. For easier comparison, we have applied heatmap visualization of these values (scale from white to red for YC3.6 signal, and from white to green for EPAC signal). NT = not tested. PSGR = prostate-specific G protein-coupled receptor. References: [1] Verbeurgt et al. (2014), [2] AktaAktaş et al. (2017), [3] Jovancevic et al. (2017b), [4] Flegel et al. (2013), [5] Giandomenico et al. (2013), [6] Maßberg et al. (2016), [7] Gelis et al. (2016), [8] Neuhaus et al. (2009), [9] Xu et al. (2000), [10] Jovancevic et al. (2017a), [11] Saito et al. (2009), [12] Adipietro et al. (2012).

| OR | expression | odorant tested | YC3.6 signal |  |  | EPAC signal |  |  |
| --- | --- | --- | --- | --- | --- | --- | --- | --- |
|  |  |  | no OR (--) | RhoOR variant | LROR variant | no OR (--) | RhoOR variant | LROR variant |
| OR2A25 | OE [1] | geraniol (5 μM) [11] | 59.8 | 62.1 | 63.7 | 60 | 61.1 | 60.6 |
| OR2J2 | OE [1], HEK293 [2] | 1-octanol (100 μM) [11, 12] | 59.1 | 59.2 | 72.4 | 61.5 | 64.7 | 64.9 |
| OR2M7 | OE [1; weak] | β-citronellol (200 μM) [11, 12] | 58.8 | 64.4 | 68 | 62.9 | 62.3 | 61.9 |
| OR5K1 | OE [1] | eugenol (10 μM) [12] | 58.1 | 68.6 | 66.4 | 60.9 | 64.5 | 66.4 |
| OR5P3 | OE [1], retina [3] | (+)-carvone (10 μM) [11] | 58 | 59.5 | 59.2 | 62.6 | 62.7 | 67.4 |
| OR8D1 | OE [1], breast, testis, sperm [4] | solotone (1 mM) [12] | 60 | 60.1 | 61.1 | 60 | 63 | 63.4 |
| OR8K3 | OE [1] | (+)-menthol (10 μM) [12] | 57.9 | 59.3 | 65.3 | 61.7 | 61.5 | 63.3 |
| OR11A1 | OE [1] | 2-ethyl fenchol (2 μM) [12] | 60.6 | 62.6 | 69.6 | 61.2 | 63.1 | 67.2 |
| OR51E1/PSGR2 | OE [1], HEK293 [2], prostate and 13 other different tissues [4], lung carcinoids [5], prostate cancer cells [6] | nonanoic acid (100 μM) [11] | 58 | 63 | 64.2 | 64.7 | 69.1 | 88.7 |
| OR51E2/PSGR | OE [1], 14 different tissues [4], melanocytes [7], prostate epithelium & prostate cancer cells [4, 8, 9], retina [10] | β-ionone (50 μM) [8] | 59.9 | 73.4 | 83.2 | 61.6 | 64.9 | 64 |
| OR51L1 | OE [1] | hexanoic acid (100 μM) [11] | 58.5 | 58.8 | 61.5 | 61.2 | 63.4 | 65.4 |
| OR1C1 | OE [1], testis [4] | linalool (1 mM) [12] | 57.9 | 67.9 | 67.2 | NT |  |  |
| OR2J2 | see above | coumarin (1 mM) [11] | 59.2 | 59.9 | 65.4 | NT |  |  |
| OR51E2/PSGR | see above | propionic acid (200 μM) [11, 12] | 59.7 | 59.4 | 62.5 | NT | | |
| -- |  | DMSO | 58 |  |  | 59,7 |  |  |

Cells were stimulated with an appropriate odorant or DMSO (control cells, see below) 38 s after the start of the measurements, and 120 s after the initial (odorant) stimulus the cell responsiveness was confirmed upon stimulation with carbachol or forskolin (see Materials & Methods). To quantify YC3.6 and EPAC signals in various conditions, we summed (individually per condition) all values in the 120 s time window (i.e. time window during which the cells are exposed to the odorant) and divided them by the number of cells measured. Such obtained values are reported in Table 1 for easier comparison.

We were concerned whether the hydrophobic character of some odorants tested could significantly delay or hamper their diffusion in aqueous environment, necessary to stimulate ORs. However, the chosen time window of 120 seconds proved sufficient, as almost all OR-mediated responses occurred without any intrinsic delay. The occasional delay present in our data (Fig. 1 and 2; Supp. Fig. 1 show the data from Fig. 2 in a manner allowing easier comparison) usually did not exceed 25 ms and could most likely be attributed to the method of odorant/reagent addition: 1 μL of odorant, carbachol or forskolin was added manually onto the sample and mixed in by pipetting. As said before, the 120 s window was sufficient to record the OR-mediated responses, although in this relatively short time the odorant-stimulated cAMP production reached a plateau only with three odorants: 1-octanol, solotone, and (+)-menthol (Supp. Fig. 1). For the remaining odorants, increased EPAC signals can be obtained with longer acquisition times.

**Figure 1.**
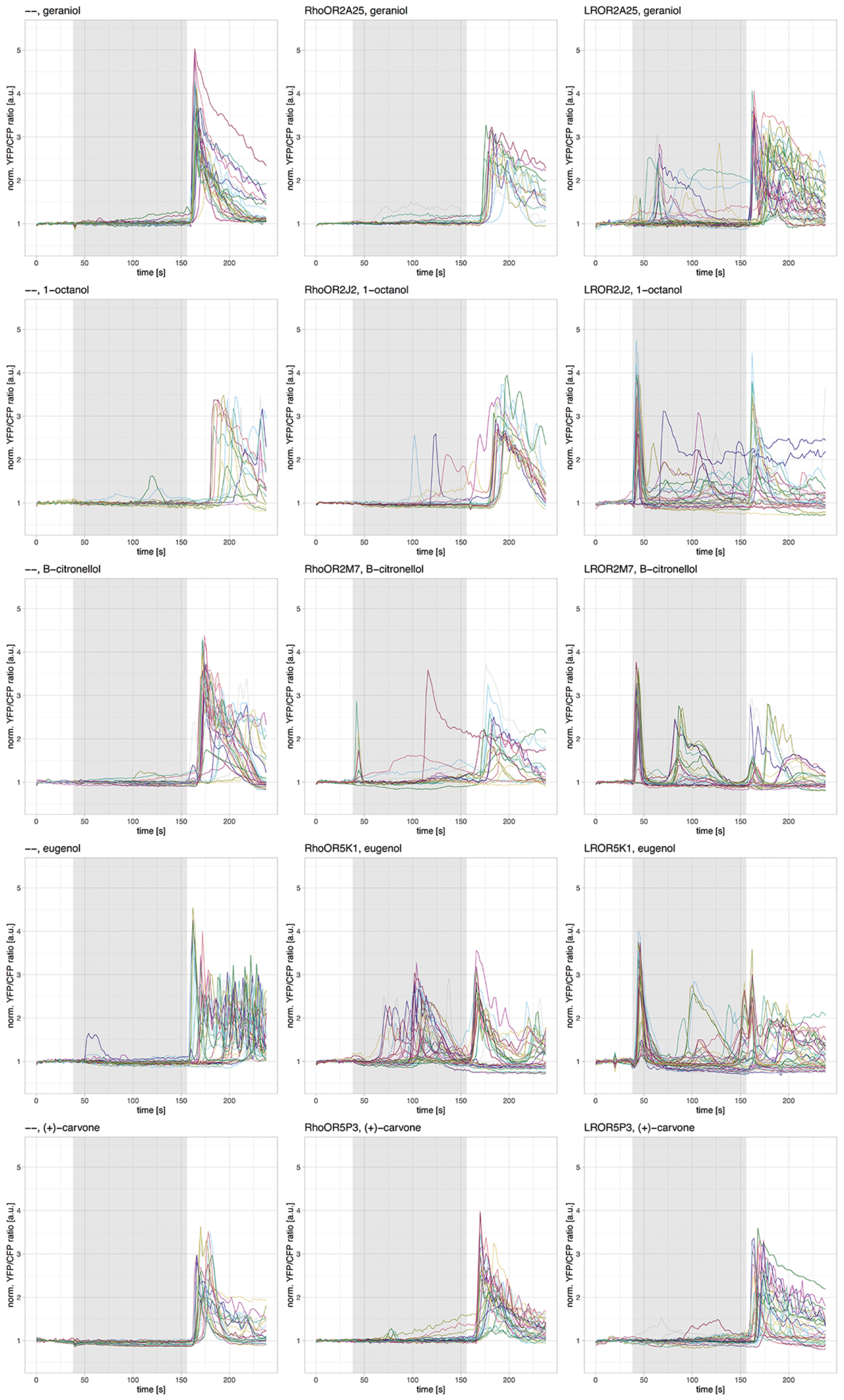
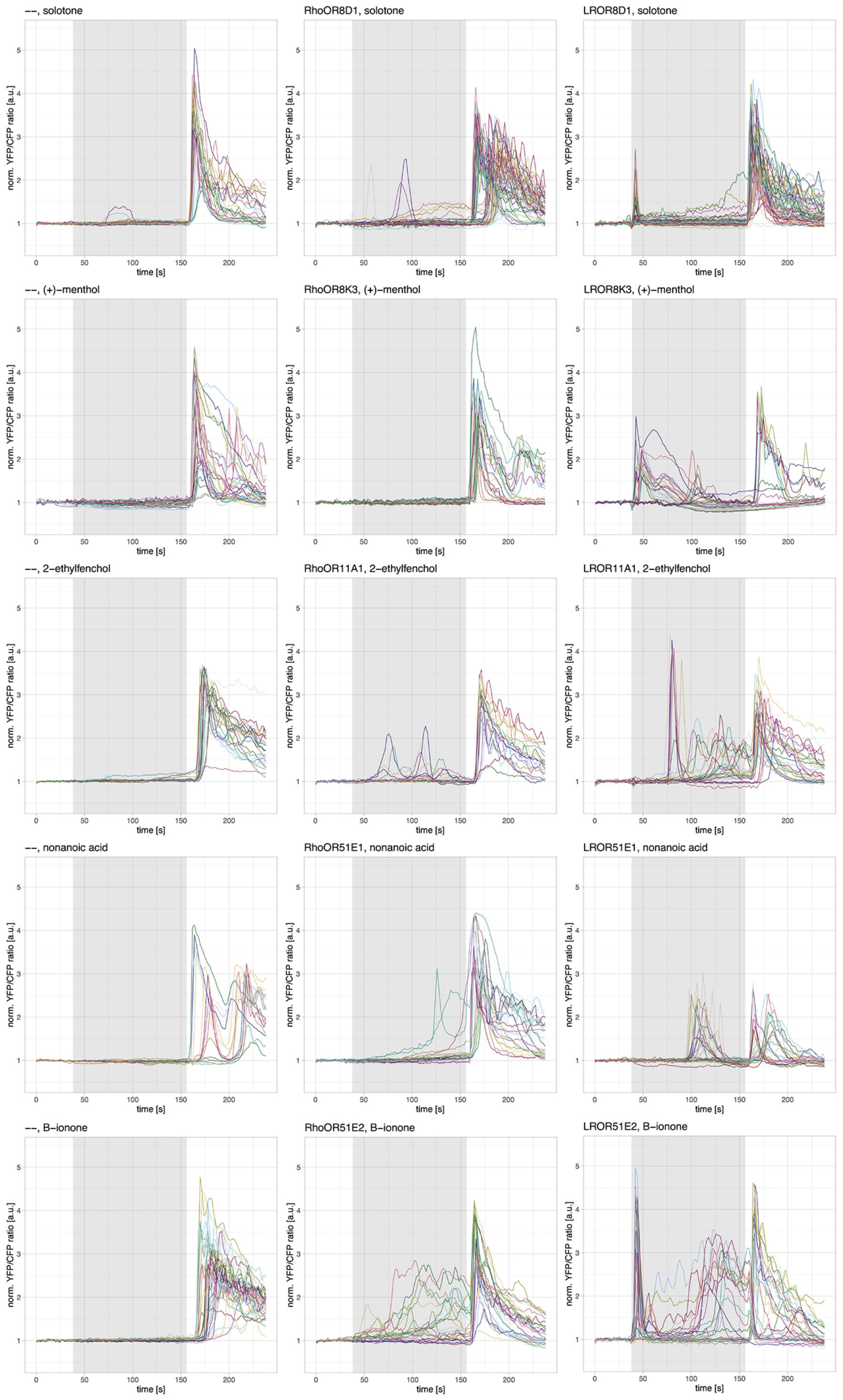
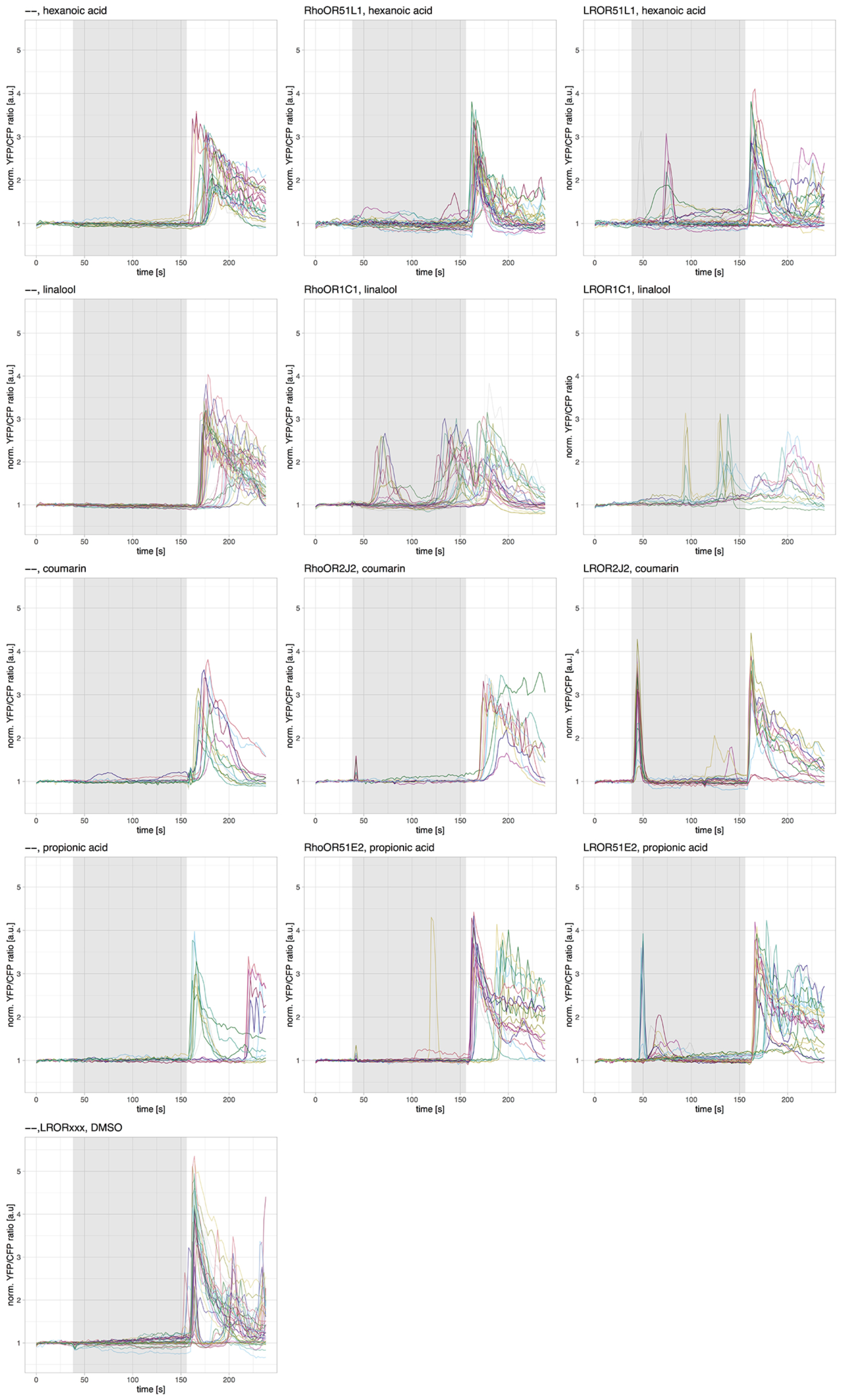
Olfactory Receptor-mediated calcium transients in HEK293TN cells. HEK293TN cells were co-transfected with RTP1s, YC3.6 and: no additional OR construct (--), RhoOR variant or LROR variant. The respective odorant was added at 38 s and 100 μM carbachol at 158 s; the grey box indicates the 120 s time window during which cells were exposed to only the odorant. Lines show the ratio change of YFP/CFP fluorescence in individual cells. All DMSO-mediated control measurements (--/LRORxxx) were pulled together and are represented in the bottom most graph.

**Figure 2.**
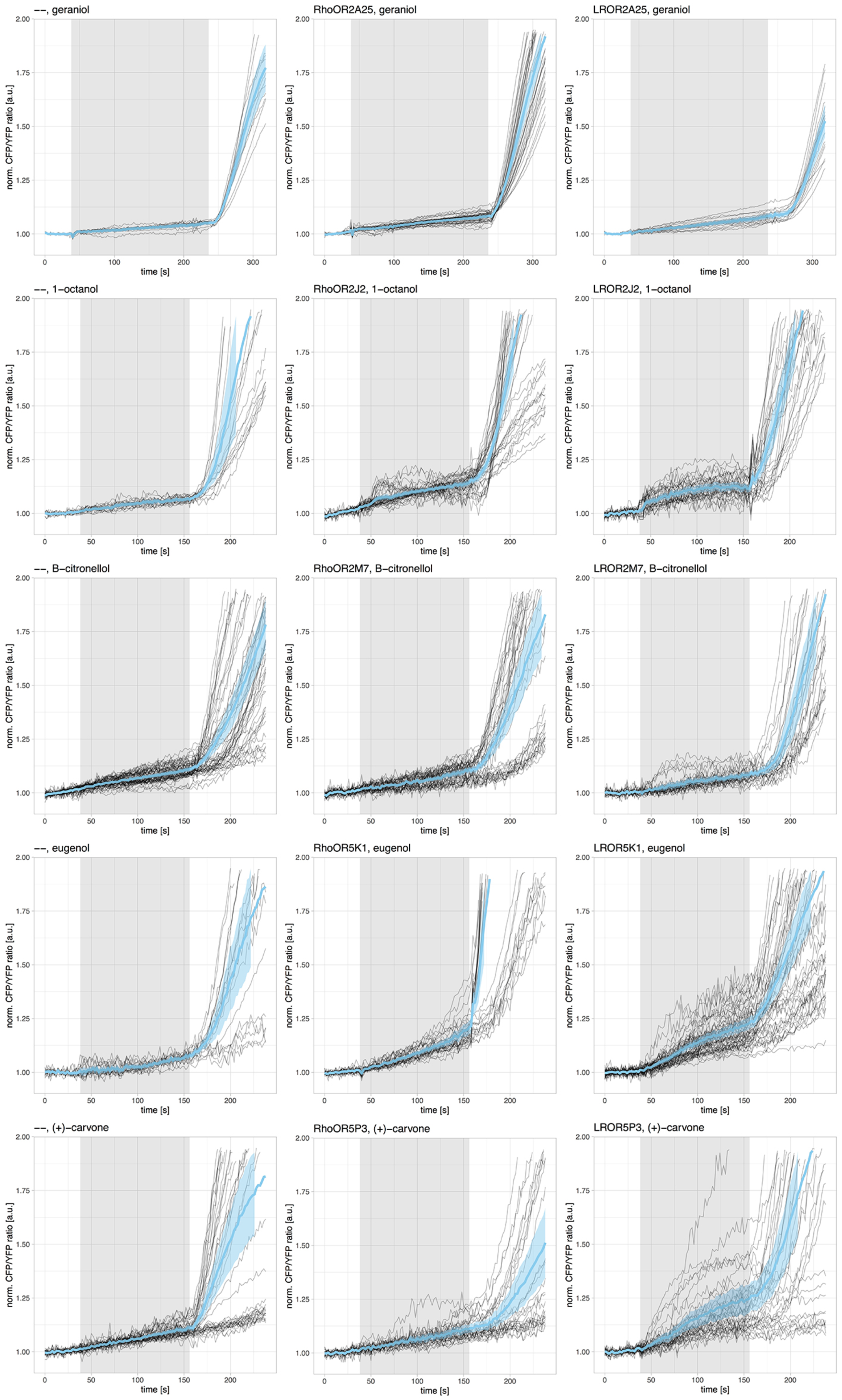
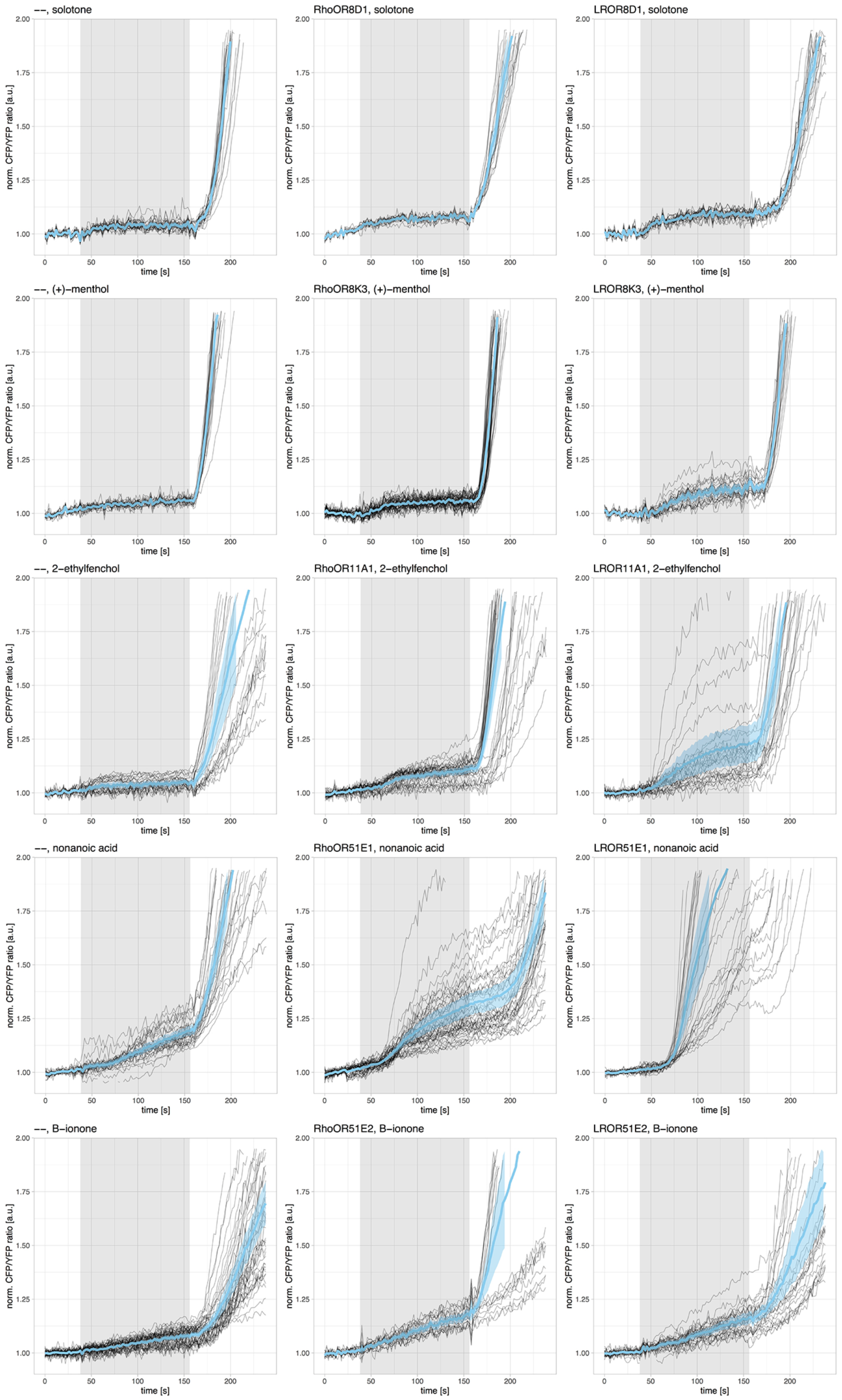
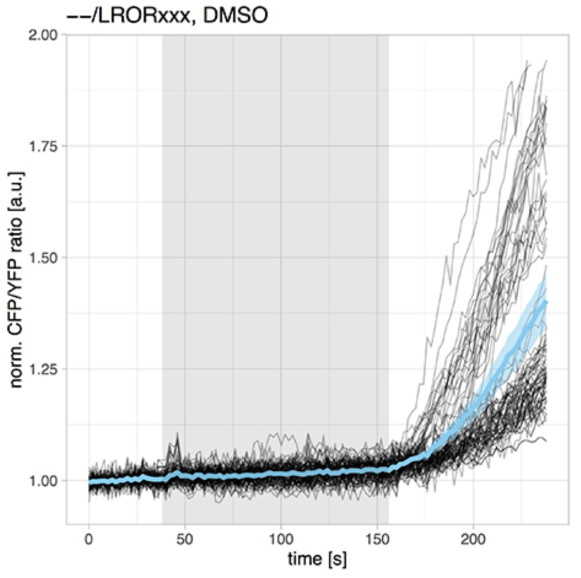
Olfactory Receptor-mediated cAMP production in HEK293TN cells. HEK293TN cells were co-transfected with RTP1s, EPAC and: no additional construct (--), RhoOR variant or LROR variant. The respective odorant was added at 38 s and 25 μM forskolin/IBMX at 158 s; the grey box indicates the 120 s time window during which cells were exposed to only the odorant. Black lines show the ratio change of CFP/YFP fluorescence in individual cells; the average ratio change (±95% confidence intervals) is shown as a thicker cyan line. All DMSO-mediated control measurements (--/LRORxxx) were pulled together and are represented in the bottom most graph.

### Many odorants trigger endogenous response in HEK293TN cells

In order to confirm that the odorant-stimulated responses are mediated by the ORs transiently expressed in HEK293TN cells, we analyzed responses of cells co-expressing only the RTP1s and YC3.6 or EPAC biosensor. As a control, we analyzed DMSO-mediated responses of: 1) HEK293TN cells co-expressing the RTP1s and biosensor, and 2) HEK293TN cells co-expressing the RTP1s, biosensor, and an LROR construct. LROR variants of all receptors were tested individually. Their DMSO-mediated responses were virtually the same as the response of cells co-expressing only the RTP1s and biosensor: no or sporadic, weak increase of intracellular Ca^2+^ level (with no obvious peaks visible in Ca^2+^ transients) (Fig. 1); 2,3±0,2% increase in cAMP production (Fig. 2). Therefore, all control measurements were pulled together (individually for YC3.6 and EPAC) and abbreviated as --/LRORxxx, DMSO (Fig. 1 and 2, bottom graphs; Supp. Fig. 1, red traces).

Interestingly, we observed endogenous response of HEK293TN cells to almost all odorants tested, with an exception of linalool (this odorant was not tested with EPAC) and eugenol (although we did measure a weak EPAC signal upon stimulation of control cells) (see Table 1 for an overview). These endogenous responses were most often observed with both biosensors, with an exception of geraniol and solotone (no endogenous response measured with EPAC), and (+)-carvone, (+)-menthol and nonanoic acid (no endogenous response measured with YC3.6) (Table 1, Fig. 1 and 2). It is important to note that two ORs (OR2J2 and OR51E2) were found to be expressed in HEK293 cells (Aktaş et al. 2017). Their presumed expression also in HEK293TN cells would explain the endogenous response observed upon stimulation of those cells with 1-octanol and coumarin (OR2J2) or ß-ionone and propionic acid (OR51E2). The remaining 10 ORs tested are not expressed in HEK293 cells (Aktaş et al. 2017), although OR1C1, OR5P3, OR8D1, and OR51E1 were found to be expressed in at least one tissue other than olfactory epithelium (see Table 1 for an overview). In total 45 ORs have been found to be expressed in HEK293 cells (Aktaş et al. 2017; see Supp. Data for an overview). HEK293 cells also express other receptors implemented in the perception of some of the chemicals tested, e.g. free fatty acid receptor 2 (FFA2) for propionic acid (Brown et al. 2003) or peroxisome proliferator-activated receptor (PPAR) α and γ for linalool, citronellol and geraniol (Katsukawa et al. 2011, Jun et al. 2014). At the moment we cannot conclude if all endogenous responses are mediated by the respective ORs or by an another (olfactory) receptor.

### LROR-variants show superior performance over the respective RhoOR constructs

We were able to record odorant-stimulated responses in real time on both biosensors for all eleven OR-odorant pairs tested (Fig. 1 and 2). This corroborates previous reports and demonstrates the feasibility of our experimental strategy. Only in case of OR2M7 were we not able to record any response above the endogenous level using the EPAC biosensor. In addition, we tested three more odorant-OR pairs only on the YC3.6 biosensor: linalool-OR1C1, coumarin-OR2J2, and propionic acid-OR51E2 (Fig. 1).

We noted that for almost all ORs tested, cells expressing the LROR variant responded more robustly: more cells responded to the odorant stimulus and the response had a higher amplitude and/or occurred faster than the response of cells expressing the respective RhoOR variants. This better performance of the LROR variants was more pronounced with the YC3.6 biosensor (Fig. 1, see Table 1 for an overview). For example, with an exception of OR1C1, OR5K1, and OR5P3, more cells expressing the LROR variants of receptors yielded odorant-stimulated Ca^2+^ transients. In case of OR8K3, we actually observed peaks in Ca^2+^ transients only with cells expressing the LROR variant of the receptor. In addition, the response of cells expressing the LROR variant had usually higher amplitude (observed for OR2A25, OR2J2, OR2M7, OR8D1, OR11A1, OR51E1, OR51E2, OR51L1) and occurred faster (observed for OR2A25, OR2J2, OR5K1, OR8D1, OR51E2) than in cells expressing the respective RhoOR variants (Fig. 1). Only in case of OR5K1 did we measure Ca^2+^ transients more robustly with the RhoOR variant.

When the activity of these ORs was monitored with EPAC, however, nearly half of them gave virtually the same responses, irrespectively of the character of their N-terminal tag (Fig. 2 and Supp. Fig. 1, see Table 1 for an overview). The more robust performance of the LROR variant was still observed for OR5K1, OR5P3, OR8K3, OR11A1, OR51E1, and OR51L1, mostly as an increased amplitude (in case of LROR51L1 the high amplitude response occurred only rarely). In case of OR5K1 the difference in amplitude was mostly observed at earlier time points, whereas at the time of forskolin addition the RhoOR and LROR variants of this receptor performed quite similarly. As it was the case with YC3.6 biosensor, we observed (+)-menthol-induced cAMP production only with LROR8K3-expressing cells, and not with the cells expressing the respective RhoOR variant.

Better performance of Lucy-Rho tagged ORs, as compared to Rho tagged variants, was previously reported by Shepard and co-workers (2013); out of 15 ORs tested, 8 ORs fused to a combined Lucy-Rho tag were expressed at the cell surface even in the absence of RTP1s, Gα_olf_, and Ric8b co-expression, as opposed to only 4 Rho tagged ORs that reached the surface without the co-expression of those accessory proteins. However, functional activity assays were performed only for two Lucy-Rho tagged ORs (Shepard et al. 2013). Here we extended this work for the analysis of additional twelve ORs. In summary, out of 14 odorant-OR pairs tested, we observed better performance of Lucy-Rho tagged ORs for five odorant-OR pairs (OR5K1, OR8K3, OR11A1, OR51E1, and OR51L1) with both biosensors, and for additional eight pairs with only one biosensor (OR2A25, 1-octanol-OR2J2, coumarin-OR2J2, OR2M7, OR8D1, ß-ionone-OR51E2, propionic acid-OR51E2 with YC3.6; OR5P3 with EPAC).

The analysis and functional characterization of ectopically expressed human ORs is becoming increasingly important as many ORs have been identified in several healthy and cancerous tissues, and show tumor-specific regulation (reviewed in Kang & Koo 2012, see also Weng et al. 2005, Cunha et al. 2006, Leja et al.2009, Cui et al. 2013, Giandomenico et al. 2013, Pronin et al. 2014, Sanz et al. 2014, 2017, Cao et al. 2015, Maßberg et al. 2015, 2016, Flegel et al. 2016, Manteniotis et al. 2016ab, Jovancevic et al. 2017b, Weber et al. 2017). Some ORs have even emerged as specific diagnostic and therapeutic targets for various cancers (Xu et al. 2000, Weng et al. 2005, Leja et al. 2009, Giandomenico et al. 2013, Muranen et al. 2011, Cui et al. 2013, Morita et al. 2016) and in the process of wound healing (Busse et al. 2014). It is foreseeable that analysis of these and other ORs in the future would require dynamic measurements of their activity.

Here we demonstrated the suitability of two biosensors routinely used for kinetic measurements of Ca^2+^ and cAMP signaling downstream of various GPCRs (Sanford & Palmer 2017) in the analysis of OR-mediated signaling. We also showed that co-expression of just one accessory protein, RTP1s, is sufficient for robust functional expression of various Lucy-Rho tagged ORs in HEK293TN cells. These minimal requirements (co-expression of LROR variant with RTP1s) could therefore be a good starting point in a situation when higher expression levels of other ORs is needed for robust analysis.

## MATERIALS and METHODS

### Constructs

pCI RhoOR2J2, RhoOR2M7, RhoOR5P3, RhoOR51E1, RhoOR51E2 and RhoOR51L1 were purchased from the addgene.org. pCI RhoOR1C1, RhoOR2A25, RhoOR5K1, RhoOR8D1, RhoOR8K3 and RhoOR11A1 were a kind gift from Hiroaki Matsunami (Duke University Medical Center, North Carolina. We did not include highly promiscuous ORs, such as OR1A1 and OR2W1, in our analyses. Rho variants of these ORs in pCI expression vector (Promega) were generated in Matsunami lab. Lucy-Rho variants were generated by swapping the Rho tag in the existing pCI RhoOR constructs for the Lucy-Rho tag (restriction cloning with NheI and EcoRI). If necessary, the respective restriction sites within the OR coding sequence were removed with silenced mutagenesis.

**Lucy**-Rho tag (NheI and EcoRI sites in italics):

**MRPQILLLLALLTLGLA**MNGTEGPNFYVPFSNATGV

*gctagc***Atgagaccccagatcctgctgctcctggccctgctgaccctaggcctggct**atgaatggcacagaaggccctaacttct acgtgcccttctccaatgcgacgggtgtggtacgc*gaattc*

Plasmid encoding the YC3.6 sensor (plasmid #67899) is available from addgene.org. The EPAC plasmid was reported before (^T^EPAC^VV^; Klarenbeek et al. 2011).

### Odorants and reagents

All odorant were purchased from Sigma Aldrich/Merck: geraniol (CAS# 106-24-1), 1-octanol (111-87-5), ß-citronellol (106-22-9), eugenol (97-53-0), (+)-carvone (2244-16-8), solotone = 4,5-dimethyl-3-hydroxy-2,5-dihydrofuran-2-one (28664-35-9), 2-ethyl fenchol (18368-91-7), nonanoic acid (112-05-0), ß -ionone (14901-07-6), hexanoic acid (142-62-1), linalool (78-70-6), coumarin (91-64-5), propionic acid (79-09-4). Odorants were prediluted in dimethyl sulfoxide (DMSO) and then diluted in the microscopy medium to the final concentration, so that the DMSO concentration did not exceed 0.1% (v/v). DMSO for control stimulations was diluted to the same 0.1% (v/v) concentration.

An adenylyl cyclase activator (25 μM Forskolin/IBMX) and 100 μM carbachol (Sigma Aldrich/Merck), activating Ca^2+^ signaling downstream from the endogenous muscarinic receptors in HEK293TN cells, served as a positive control.

### Cell culture & sample preparation

HEK293TN cells (System Biosciences, LV900A-1) were cultured using Dulbecco’s Modified Eagle Medium (DMEM) supplied with glutamax, 10% FBS, penicillin (100 U/ml) and streptomycin (100 ug/ml), and incubated at 37°C and 5% CO_2_. All cell culture regents were obtained from Invitrogen (Bleiswijk, NL). Cells were transfected in a 35 mm dish holding a glass coverslip (24 mm ∅, Menzel-Gläser, Braunschweig, Germany), using polyethylenimine (3 μL of PEI:1 μL of DNA) according to the manufacturer’s protocol. For each transfection, we used 500 ng of OR-carrying plasmid, 250 ng of RTP1s-carrying plasmid, and 200-300 ng of YC3.6 or EPAC-carrying plasmid, respectively. Rho tagged- and Lucy-Rho tagged ORs were co-expressed in HEK293TN cells with either RTP1s or RTP1s fusion to a fluorescent protein, mCherry (we observed no differences in the ability to support functional expression of ORs between RTP1s and mCherry-RTP1s fusion; data not shown). Samples were imaged one day after transfection: coverslips were mounted in an Attofluor cell chamber (Invitrogen, Breda, NL) and submerged in 1 mL microscopy medium (20 mM HEPES, pH=7.4, 137 mM NaCl, 5.4 mL KCl, 1.8 mM CaCl_2_, 0.8 mM MgCl_2_, and 20 mM glucose; Sigma Aldrich/Merck). All experiments were performed at 37°C.

### Wide-field microscopy

Ratiometric FRET measurements were performed on a previously described wide-field microscope (van Unen et al. 2015). Typical exposure time was 100 ms, and camera binning was set to 4×4. Fluorophores were excited with 420/30 nm light and reflected onto the sample by a 455DCLP dichroic mirror. CFP emission was detected with a BP470/30 filter and YFP emission was detected with a BP535/30 filter by rotating the filter wheel. Before each FRET acquisition series, mCh-RTP1s expression in cells was confirmed. To this end, RFP was excited with 570/10 nm light reflected onto the sample by a 585 dichroic mirror. RFP emission was detected with a BP620/60 nm emission filter. Simultaneous co-expression of the RhoOR- or LROR-constructs in these cells could not be verified. All acquisitions were corrected for background signal and bleedthrough of CFP emission in the YFP channel (55% of the intensity measured in the CFP channel).

### Image and statistic analyses

ImageJ (National Institute of Health) was used to analyze the raw microscopy images. Background subtractions, bleedthrough correction and calculation of the normalized ratio per time point per cell were done in Excel (Microsoft Office). All plots were prepared with the PlotTwist web app (Goedhart 2019). Plots show the average response as a thicker line and a ribbon for the 95% confidence interval around the mean.

## Author contributions

APB designed and performed experiments, analyzed the data, and wrote the manuscript. LvW provided technical assistance with cell culture maintenance. JG participated in study design, data interpretation, and data visualization. All authors approved the final manuscript.

## Acknowledgments & Funding

We thank Hiroaki Matsunami (Duke University Medical Center, North Carolina) for providing the constructs. A Pietraszewska-Bogiel was supported by NanoNextNL, a micro and nanotechnology consortium of the Government of the Netherlands and 130 partners.

## SUPPLEMENTARY DATA

**Supp. Figure 1.**
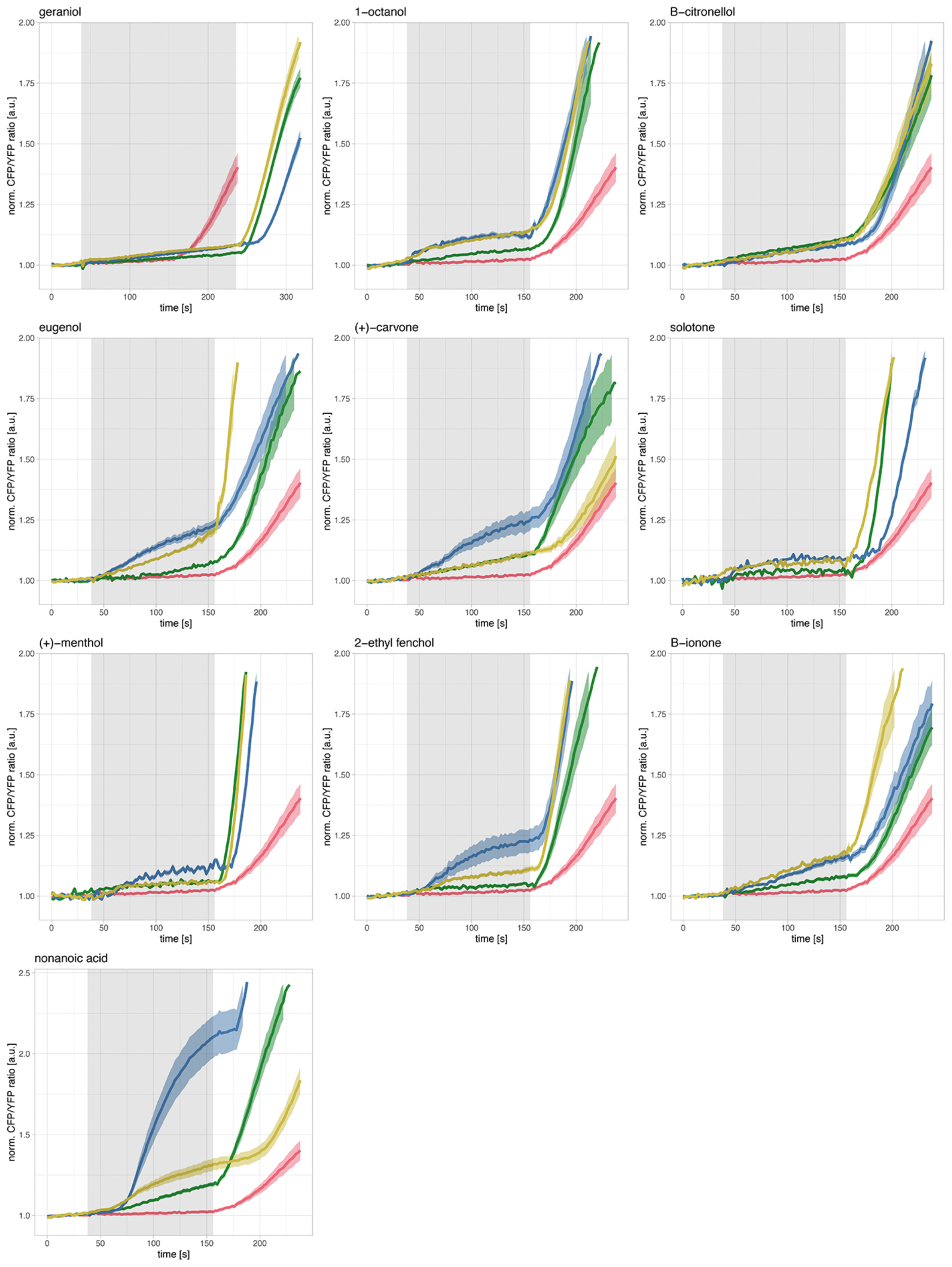
Odorant-mediated cAMP production in HEK293TN cells. HEK293TN cells were co-transfected with RTP1s, EPAC and: no additional OR construct (--, yellow traces), the respective RhoOR variant (green traces) or the respective LROR variant (blue traces). All DMSO-mediated responses measured in control cells (--/LRORxxx) were pulled together and are represented with red traces. The respective odorant (specified in the graph title) or DMSO was added at 38 s and 25 μM forskolin/IBMX at 158 s (please note later addition of forskolin/IBMX [at 238 s] in the top left graph, which does not apply to the DMSO control trace). All time traces show the average ratio change of CFP/YFP fluorescence (±95% confidence intervals).

Olfactory receptors expressed in HEK293 cells, as reported by Aktaş and colleagues (2017) (deorphanized ORs are highlighted in bold): OR1F1, OR1J1, OR1K1, OR2A4, OR2AG1, OR2AG2, OR2AK2, OR2B2, OR2B6, **OR2C1**, OR2D2, OR2D3, OR2H2, **OR2J2**, OR2W3, OR4C3, OR4F3, OR4F16, OR4F29, OR4K2, OR5C1, OR5H1, OR6A2, OR6B2, OR6V1, OR7D2, OR7E24, OR9A2, OR9A4, OR10A2, OR10A4, OR10A5, OR10AD1, OR10C1, OR11H2, OR11H12, OR12D3, OR13A1, OR13J1, **OR51E2**, OR52E2, OR52B6, OR52I2, OR52W1, OR56B4.

In addition, HEK293 cells also express 10 pseudogenes: OR1F2P, OR6W1P, OR7E2P, OR7E5P, OR7E12P, OR7E14P, OR7E37P, OR7E91P, OR7E156P, OR10V2P.

